# The amount of nitrogen used for photosynthesis governs molecular evolution in plants

**DOI:** 10.1101/162693

**Authors:** Steven Kelly

## Abstract

**Background:** Genome and transcript sequences are composed of long strings of nucleotide monomers (A, C, G and T/U) that require different quantities of nitrogen atoms for biosynthesis.

**Results:** Here it is shown that the strength of selection acting on transcript nitrogen content is determined by the amount of nitrogen plants require to conduct photosynthesis. Specifically, plants that require more nitrogen to conduct photosynthesis experience stronger selection on transcript sequences to use synonymous codons that cost less nitrogen to biosynthesise. It is further shown that the strength of selection acting on transcript nitrogen cost constrains molecular sequence evolution such that genes experiencing stronger selection evolve at a slower rate.

**Conclusions:** Together these findings reveal that the plant molecular clock is set by photosynthetic efficiency, and provide a mechanistic explanation for changes in plant speciation rates that occur concomitant with improvements in photosynthetic efficiency and changes in the environment such as light, temperature, and atmospheric CO_2_ concentration.

## Introduction

Cells are built from macromolecules (proteins, RNA, DNA, phospholipids and polysaccharides) that in turn are constructed from monomers (amino acids, nucleotides, fatty acids and sugars). The majority of plants can biosynthesise all of the monomers and macromolecules they require from inorganic carbon (CO_2_) and nitrogen (NO_3_/NH_4_) obtained from their environment. Of these two resources, nitrogen is scarcer and hence plant growth rate is generally nitrogen limited in both natural and agricultural environments^1–4^. This limitation in growth is caused by the fact that synthesis of proteins required for photosynthetic carbon assimilation needs a substantial nitrogen investment^5,6^. Therefore, the rate of carbon acquisition and macromolecule biosynthesis (and hence growth) in plants is predominantly limited by nitrogen availability^6^.

Photosynthetic nitrogen use efficiency (PNUE) is the amount of carbon that can be fixed per unit of nitrogen invested by the plant. Multiple disparate anatomical, physiological and molecular factors contribute to variation in PNUE such that there is a large variation between different plant species^7^. For example, plants that use the C_4_ photosynthetic pathway exhibit higher nitrogen use efficiency when compare to plants that use C_3_ photosynthesis. The cohort of changes that facilitated C_4_ evolution enabled plants to reduce resource allocation to photosynthetic machinery without causing a corresponding reduction in photosynthetic rate^8^. Thus, C_4_ plants can achieve ~50% higher rates of photosynthesis than C_3_ plants given the same amount of nitrogen^9^.

Nucleotide monomers (A, C, G and T/U) differ in their biosynthesis requirements, with different nucleotides requiring different quantities of nitrogen atoms for their construction. Adenine and guanine require 5 nitrogen atoms for their biosynthesis, cytosine requires 3, and thyamine/uracil only require 2. While the sequence and abundance of proteins within a cell are functionally constrained, it is possible to encode the same polypeptide with multiple different nucleotide sequences by using different synonymous codons. This redundancy in the genetic code, coupled with the difference in nucleotide nitrogen content, means that it is possible to reduce the allocation of cellular resources to transcript sequences without altering protein sequence or function^10^. For example, a single A to T substitution in a highly expressed transcript such as RuBisCo small subunit (~5000 transcripts per cell) saves an equivalent amount of nitrogen (~15000 atoms) as is contained in three complete RuBisCo holoenzymes (~5000 nitrogen atoms per hexadecamer). It was thus hypothesised that plants that require increased quantities of nitrogen to fix CO_2_ would be more nitrogen limited and thus natural selection would favour codons in transcript sequences that required less nitrogen to biosynthesise.

Here it is shown that plants that require more nitrogen to conduct photosynthesis experience stronger selection to minimise transcript nitrogen biosynthesis cost. It is also shown that these plants exhibit a stronger mutation bias away from GC and thus have lower genome-wide GC content. Furthermore, it is demonstrated that the strength of selection acting on transcript sequence biosynthesis cost explains a significant proportion of variation in gene evolutionary rate, whereby genes that experience stronger selection to minimise cost are evolving slower than genes that experience weaker selection. Together these findings directly link the photosynthetic efficiency to plant to the rate at which its genes and genome evolve, and provide a mechanistic link between fluctuation in rates of plant diversification and changing environmental conditions.

## Results

### Variation in photosynthetic nitrogen use efficiency causes a concomitant variation in the strength of selection acting on transcript nitrogen cost

To test the hypothesis that variation in the amount of nitrogen used for photosynthesis influences the strength of selection acting on transcript biosynthesis cost, an analysis of molecular sequence evolution was conducted for 11 plant species for which both whole genome sequences^11^ and accurate photosynthetic nitrogen use efficiencies^7^ were available. This set of species included both C_3_ and C_4_ grasses, as well as C_3_ herbs and trees (Fig. 1a, Supplemental Table S1, and Supplemental File S1). For each species, the strength of selection acting on transcript nitrogen cost [*S*_*c*_] was inferred from the complete set of open reading frames in the respective genome using the SK model^12^ implemented using CodonMuSe^10^. Consistent with the hypothesis, those species that required more nitrogen to conduct photosynthesis exhibited stronger selection (more negative value for *S*_*c*_) to minimise the nitrogen biosynthesis cost of transcript sequences (R^2^ = 0.62, p < 0.001, Fig. 1b). Although, both PNUE and *Sc* exhibited significant phylogenetic signal (Supplemental File S2), correction for this signal did not remove the significant, strong positive association between these two traits (R^2^ = 0.78, p = 0.004, Supplemental File S2) and thus the amount of nitrogen used for photosynthesis in a plant determines the strength of selection acting on its transcript nitrogen biosynthesis cost.

**Figure 1.**
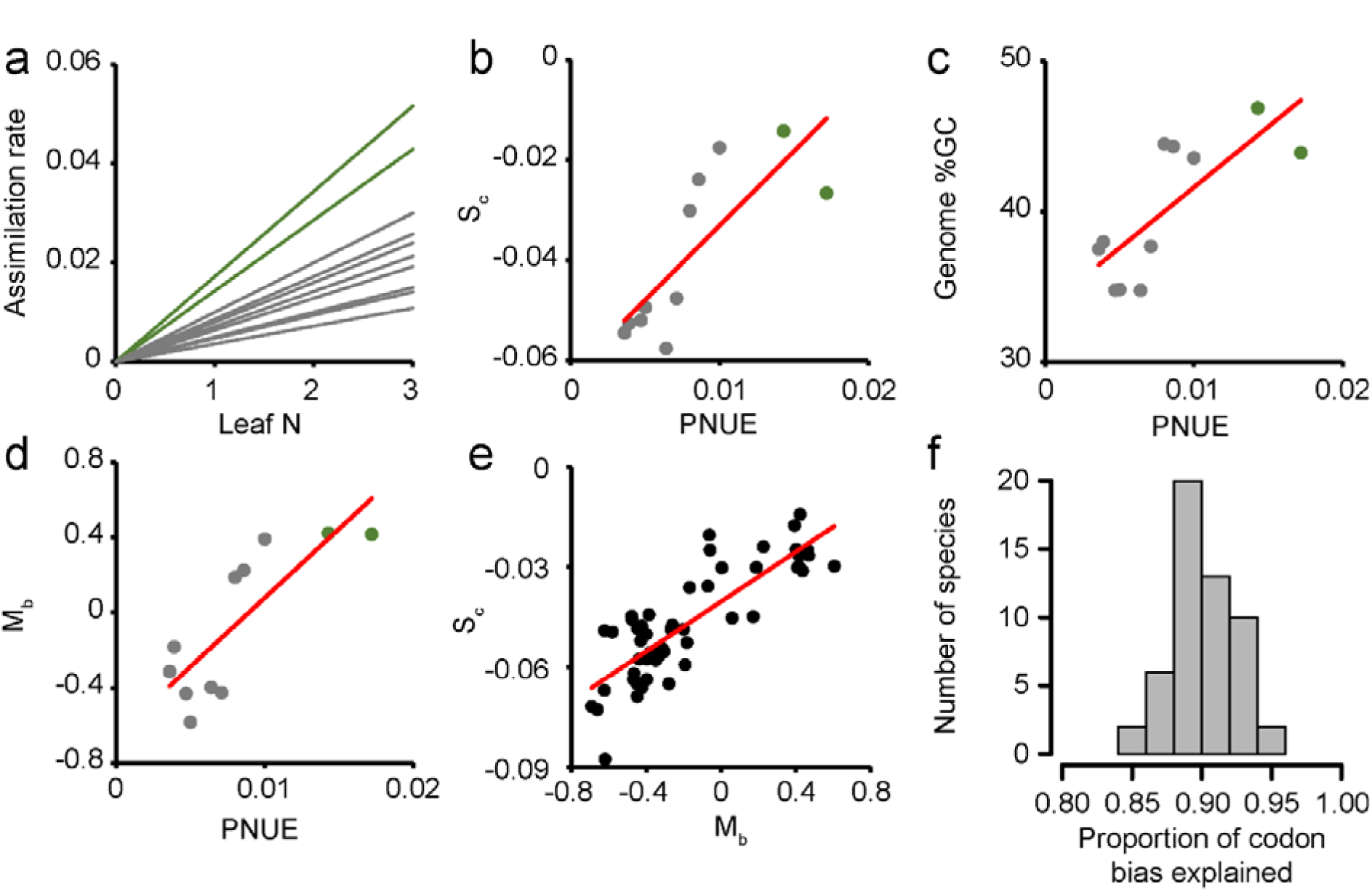
Photosynthetic nitrogen use efficiency (PNUE) drives selection acting on transcript biosynthesis cost and mutation bias. Plots in parts a to e depict the same species set. a) The range of relationships observed between light saturated CO_2_ assimilation rate (µ mol CO_2_ m^−2^ s^−1^ µ mol PAR^−1^) and leaf nitrogen (g N m^−2^). C_4_ species shown in green and C_3_ species shown in grey. Complete datasets provided in Supplemental file 1. The slope of the line is the PNUE for that species. b) The relationship between PNUE and the strength of selection acting on codon biosynthesis cost (*S*_*c*_, R^2^ = 0.62) for these species. c) The relationship between PNUE and the Genome wide GC content (R^2^ = 0.58). d) The relationship between PNUE and the strength of mutation bias acting on coding sequences (*M*_*b*_, R^2^ = 0.65). e) The relationship between *S*_*c*_ and *M*_*b*_ for all angiosperm species in Phytozome (R^2^ = 0.69). f) The proportion of codon bias that can be explained by the joint effects of mutation bias and selection acting on nitrogen biosynthesis cost.

### Variation in photosynthetic nitrogen use efficiency causes a concomitant variation in mutation bias and genome-wide GC content

As the nitrogen biosynthesis cost of DNA sequences varies (AT pairs require 7 and GC pairs require 8 nitrogen atoms), it was further hypothesised that those species that required more nitrogen to conduct photosynthesis would exhibit a stronger genome-wide mutation bias towards AT base pairs. Consistent with the hypothesis, those species that required more nitrogen to conduct photosynthesis had lower genome-wide GC content and thus invested less nitrogen in their genome sequences (R^2^ = 0.58, p < 0.001, Fig. 1c). This mutation-driven phenomenon was also apparent from the analysis of coding sequences, where codon mutation bias towards AT rich codons was stronger in species that had lower photosynthetic nitrogen use efficiencies (R^2^ = 0.65, p < 0.001, Fig. 1d). Like for *S_c_,* both genome-wide GC content and mutation bias exhibited strong phylogenetic signal (Supplemental File S2). However, in contrast to the case for *S*_*c*_ correction for phylogenetic signal reduced the strength of the positive association with PNUE such that they failed to achieve statistical significance (p ≥ 0.05, Supplemental File S2).

To exclude the possibility that low sample size caused the statistical test to fail, an additional analysis on a larger species set was conducted. If PNUE influences genome-wide GC content and mutation bias, then there should be a linear dependency between *S*_*c*_ and these traits across the angiosperm phylogenetic tree. However, if there is no association between GC content, mutation bias and PNUE then *S*_*c*_ will also be independent of GC content and mutation bias. To investigate this, a larger set of C_3_ angiosperm genomes on Phytozome were analysed to determine whether there was a global, significant, positive association between *S*_*c*_ and GC content and mutation bias. As postulated, those species that exhibited stronger selection acting on transcript biosynthesis cost also exhibited lower genome-wide GC content (R^2^ = 0.69, Fig. 1e, Supplemental Table S1). Correcting for phylogenetic signal did not remove the strong positive association (R^2^ = 0.21, *p* = 0.007, Supplemental File S2). Therefore, the most parsimonious explanation is that variance in PNUE is a major determinant of biased patterns of nucleotide use in both genome and transcriptome sequences. Moreover, when mutation bias and selection acting on transcript nitrogen cost are considered together, they are sufficient to explain ~90% of variance in genome-wide patterns of synonymous codon use in plants (Fig. 1e, Supplemental Table 1, Supplemental File S3).

### Variation in the strength of selection acting on nitrogen biosynthesis cost contributes to variation in gene evolutionary rate

Given that variance in PNUE governs the strength of selection acting on gene sequences, it was postulated that this would cause a concomitant variance in molecular evolutionary rate of genes. Specifically, those genes that experience stronger selection to minimise transcript nitrogen cost would have lower rates of molecular evolution when compared to genes that experience weaker selection. This phenomenon occurs because spontaneous mutations that increase transcript biosynthesis cost will be more deleterious in genes that experience stronger selection to minimise cost irrespective of whether that mutation is synonymous or non-synonymous^10^. As mutations that are more deleterious will be lost more rapidly, this results in a lower molecular evolution rate for genes that experience stronger selection^10^. This phenomenon has previously been observed for bacterial genes^10^.

To investigate this, both the number of synonymous substitutions per synonymous site (*K*_*s*_) and the number of non-synonymous substitutions per non-synonymous site (*K*_*a*_) were estimated from pairwise alignments of single copy orthologous genes in a set of 38 plant species (1406 pair-wise comparisons). The strength of selection acting on transcript nitrogen cost was also inferred for each individual gene^10^. For each species pair, this data was subject to multiple regression analysis to estimate the proportion of variance in *K*_*a*_ or *K*_*s*_ that was explained by variance in *S*_*c*_ between that species pair (Supplemental File S2, Supplemental Table S2). Consistent with the hypothesis, genes experienced stronger selection to minimise transcript nitrogen cost evolved more slowly than those that experience weaker selection (Figure 2, Supplemental File S2). Moreover, variance in the strength of selection explained up to 10% of variance in synonymous site evolutionary rate (Supplemental File S2) and ~2% of variance in non-synonymous site evolutionary rate across all species (Figure 2, Supplemental File S2). Thus, as genome-wide values for *S*_*c*_ are determined by PNUE, it follows that the tempo of the plant molecular clock is modulated by changes in PNUE.

**Figure 2.**
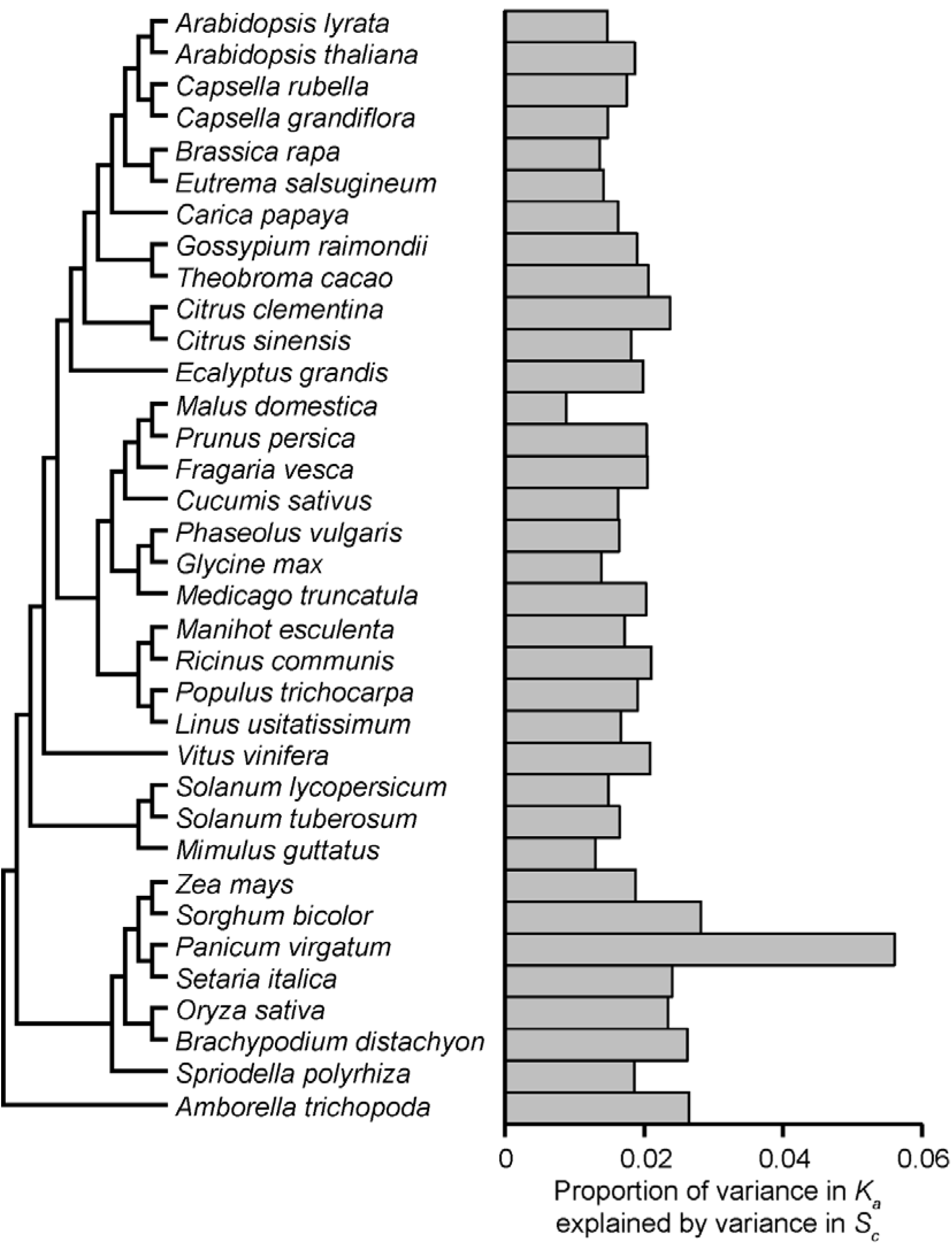
The strength of selection acting on transcript biosynthesis cost constrains the rate of amino acid sequence evolution. Grey bars depict mean values estimated from all possible pairwise comparisons featuring the species under consideration. Phylogenetic tree adapted from Phytozome^11^.

## Discussion

There is substantial inter-species variation in the amount of nitrogen required to conduct photosynthesis in plants^6,7^. In this work, it is shown that this variation is a major determinant of plant gene and genome composition, and modulates the rate at which plant gene sequences evolve. The findings presented here provide significant new insight into the relationship between metabolism, the environment and molecular evolution in plants. They are also compatible with previous reports that revealed that wild plants contained less nitrogen in their DNA when compared to domesticated relatives that had been supplemented with nitrogen fertiliser for thousands for years^13^.

Speciation and extinction rates in plants are a function of molecular substitution rate, such that lineages with higher rates of molecular substitution have higher rates of speciation and extinction^14^. Therefore, the mechanistic link between PNUE and molecular evolution presented here has significant implications for our understanding of the past, present and future of plant evolution. For example, plants with enhanced PNUE, and thus with higher rates of molecular evolution, will have therefore higher rates of speciation and extinction. As a corollary, evolutionary adaptations that increase PNUE will also increase rates of speciation and extinction. For example, the suite of molecular and anatomical changes that facilitate the evolution C_4_ photosynthesis result in a dramatic reduction in the amount of nitrogen required to conduct photosynthesis. The findings presented here predict that this increase in PNUE would cause a concomitant reduction in the strength of selection gene sequences and therefore result in an increased rate of molecular evolution. This PNUE-driven increase in molecular evolution rate provides a simple mechanistic explanation for the increase in rates of speciation that are observed concomitant with the evolution of C_4_ photosynthesis^15^.

Increases in atmospheric CO_2_ concentration cause corresponding increases in PNUE in plants. In the short term, this increase in PNUE is caused by a reduction in the rate of photorespiration^16^. In the long term, plants also adapt to higher CO_2_ concentration by reduction in the investment of cellular resources in photosynthesis protein production^17^. Thus, when atmospheric CO_2_ increases, PNUE increases. The link between PNUE and molecular evolution presented here predicts that this increase in PNUE will cause a corresponding increase in molecular evolution rate, and thus an increase in the rate of plant diversification (Figure 3A & B). This therefore provides a mechanistic explanation for the observed relationship between plant diversification rates and changes in atmospheric CO_2_ concentration^18^.

**Figure 3.**
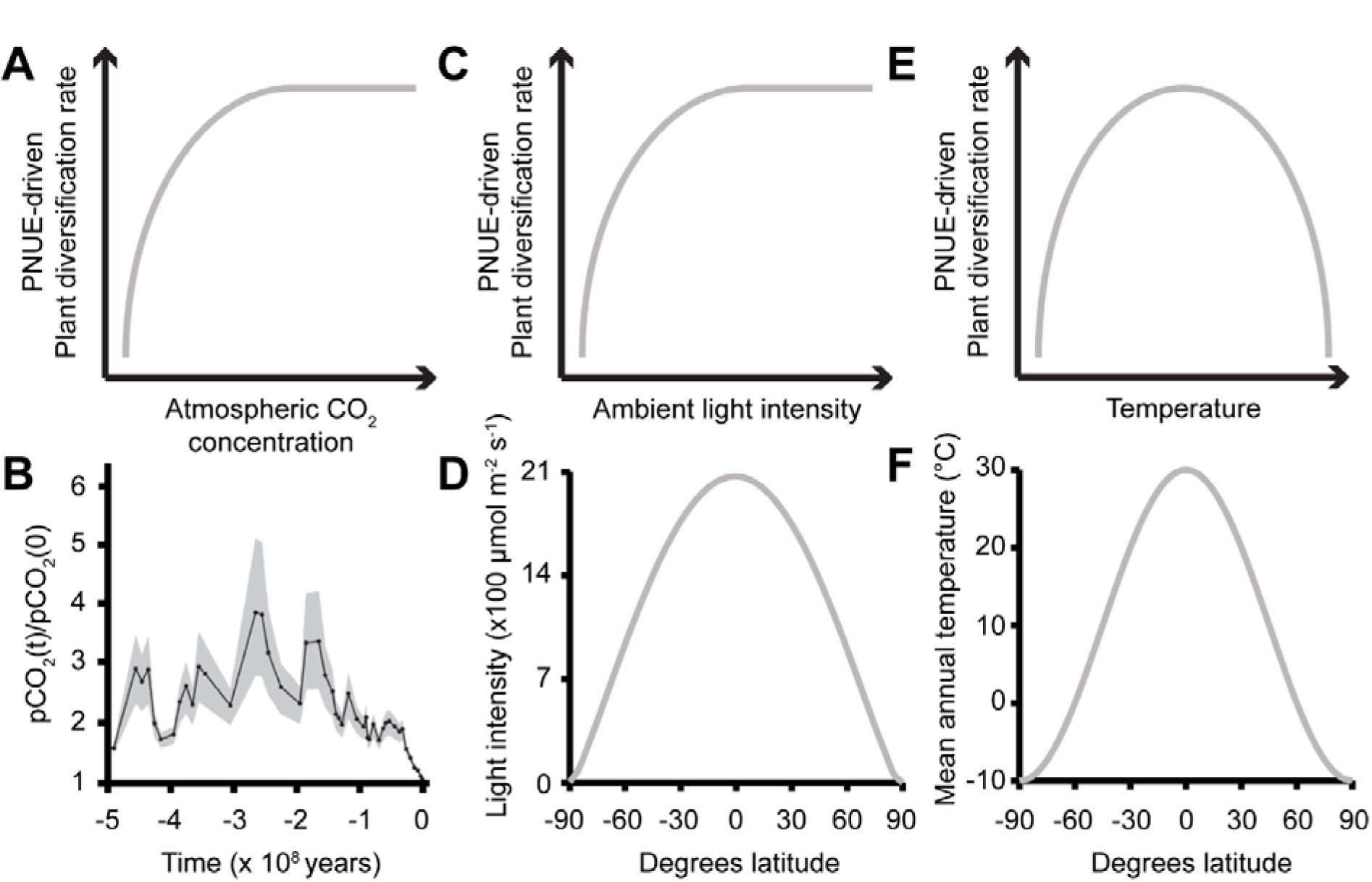
Photosynthetic nitrogen use efficiency (PNUE) drives plant diversification. A) Cartoon depicting the proposed relationship between PNUE-driven plant diversification and atmospheric CO_2_ concentration. B) Changes in partial pressure of atmospheric CO_2_ (pCO_2_) over the last 500 million years, mean indicated as solid line with grey shading for upper and lower limits adapted from^27^. C) Cartoon depicting the relationship between PNUE and light intensity. D) The photon flux density on the surface of the earth at 0° longitude at noon on the vernal equinox. E) Cartoon depicting the proposed relationship between PNUE-driven plant diversification and ambient temperature. F) The average (contemporary) annual temperature at the surface of the earth^28^.

Similar to changes in CO_2_ availability, changes in other environmental factors such as light availability (Figure 3C & D) and temperature (Figure 3E & F) also influence photosynthetic rate and thus PNUE. Unlike CO_2,_ these other environmental factors vary widely over the surface of the planet. For example, light intensity and temperature are not uniformly distributed on the surface of the earth, but instead decrease as a function of distance from the equator (Figure 3D & F). This variation is due to the curvature of the earth and the corresponding increase in the angle of the incident light. The findings presented here predict that plant diversification rates will be higher towards the equator where light and temperature are less limiting on photosynthesis and thus PNUE will be higher. These findings therefore provide additional insight into the plant species latitude diversity gradient^19,20^, where rates of plant diversification are higher in regions that are closer to the equator.

## Conclusion

In summary, plants build their genes and genomes from monomers assembled from inorganic carbon and nitrogen. Of these two, nitrogen is more limiting such that plants that require higher quantities of nitrogen to conduct photosynthesis have less nitrogen available for other uses and thus experience stronger selection to reduce nitrogen investment in gene sequences. A multitude of environmental factors can exacerbate or ameliorate PNUE. Therefore, both the environment and genetic factors can modulate the strength of selection acting to reduce nitrogen investment in gene sequences and hence modulate plant genome composition and molecular evolution. Hence, at multiple scales plant evolution is governed by the amount of nitrogen required to conduct photosynthesis.

## Methods

### Data sources

The genome sequences and corresponding set of representative gene models for each species were downloaded from Phytozome^11^. The *Helianthus annuus* genome was obtained from^21^. Photosynthetic measurements and leaf nitrogen measurements were obtained from^7^.

### Inference of selection acting on codon usage bias

To obtain the number of tRNA genes in each genome, tRNAscan^12^ was run on each of the plant genomes. For each species the tRNAscan output file and the complete set of representative coding sequences was analysed using CodonMuSe^10^. The values for mutation bias (M_b_) as well as the composite parameters of selection acting on transcript biosynthesis cost (*S*_*c*_), and selection acting on translational efficiency (*S*_*t*_) were obtained (Supplemental Table S1). CodonMuSe by default estimates the proportion of variance in codon use that can be explained by the mutation bias and selective forces.

### Phylogenetic tree inference for the 11 species with PNUE data

The complete set of proteomes for the 11 species used in this analysis were subject to orthogroup inference using OrthoFinder^22^. In the case of hexaploid wheat genome, only proteins derived from genes present in the wheat A genome were used for orthogroup inference. Orthogroups containing proteins derived from single copy genes in each of the 11 species were selected and aligned using the MAFFT^23^ L-INS-i algorithm. These alignments were trimmed to remove any columns containing gap characters and then concatenated to form a multiple sequence alignment containing 4949 aligned amino acid positions in each species. This alignment and was subject to bootstrapped maximum likelihood phylogenetic tree inference using IQ-TREE^24^ while estimating the best fitting model of sequence evolution from the data. The best fitting model was inferred to be JTTDCMut+F+G4 by Bayesian information criterion. This tree was used for the phylogenetic least squares analysis and is provided in Supplemental File S2.

### *K_a_ and K*_*s*_ estimation and comparison with S_c_

The predicted proteins from 38 species were downloaded from Phytozome. These species were subject to orthogroup and ortholog inference using OrthoFinder^22^. All 1406 pairwise comparisons between species were subsequently conducted. Each pairwise comparison comprised the following steps. 1) The full set of single copy orthologs for the species pair under consideration were isolated. 2) The protein sequences for each orthologous pair were aligned using MAFFT^25^ L-INS-i and the coding sequences re-threaded back through the protein sequence alignment. 3) The resulting coding sequence alignments were parsed to remove any gap-containing columns. 4) Ungapped alignments containing more than 100 aligned codons were subject to *K*_*a*_ and *K*_*s*_ inference using KaKsCalculator v2.0^26^ using the default settings. Additional data filtering and quality control were carried out as described in Supplemental File 2. Individual estimates for *S*_*c*_ and *S*_*t*_ were obtained for each gene in the 38 species using CodonMuse. Here, the value for *M*_*b*_ in each inference was set to the genome-wide value estimated from an analysis of all genes. Pairwise species comparisons that had > 100 genes satisfying all filtration criteria were selected for further analysis.

The value for *K*_*a*_ and *K*_*s*_ are dependent on several factors

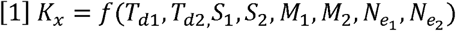

Where *K*_*x*_ is either *K*_*a*_ or *K*_*s*_,,*T*_*d1*_ is the divergence time in number of generations between species 1 and the most recent common ancestor of the species pair being analysed, *S*_*1*_ is the strength of selection acting on the sequence of the gene in species 1, *M*_*1*_ is the mutation rate species 1, and *N*_*e1*_ is the effective population size of species 1. *S*_*c*_ is a composite parameter^12^ that is a product of a component of the selection coefficient *S*_*1*_ and the effective population size *N*_*e1*_. Thus, each pairwise species comparison was subject to multiple regression analysis using the lm function in R using the following model

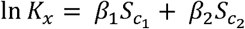

Where β_*1*_ thus incorporates both *T*_*d1*_ and *M_1_*. Thus, the multiple regression evaluates the component of variance in *K*_*a*_ or *K*_*s*_ that is attributable to both *S*_*c1*_ and *S_c2_.* The natural log of the *K_a_ and K*_*s*_ estimates were taken as both *K*_*a*_ and *K*_*s*_ are log-normally distributed whereas *S*_*c*_ is normally distributed. All data was confirmed to be normally distributed by the Shapiro-Wilks test for normality prior to use in regression analysis. The mean of the adjusted R-squared for all pairwise comparisons featuring a given species was taken as an estimate the proportion of variance that is explained by variation in *S*_*c*_ for that species.

## Declarations

Ethics approval and consent to participate

Not applicable

### Consent for publication

Not applicable

### Availability of data and material

The datasets generated and/or analysed during the current study are all provided in the supplemental material.

**Competing interests**

The author declares that he has no competing interests.

## Authors' contributions

SK conducted the study and wrote the manuscript.

## Acknowledgements

Not applicable

